# Efficient inference for agent-based models of real-world phenomena

**DOI:** 10.1101/2021.10.04.462980

**Authors:** Andreas Christ Sølvsten Jørgensen, Atiyo Ghosh, Marc Sturrock, Vahid Shahrezaei

## Abstract

The modelling of many real-world problems relies on computationally heavy simulations. Since statistical inference rests on repeated simulations to sample the parameter space, the high computational expense of these simulations can become a stumbling block. In this paper, we compare two ways to mitigate this issue based on machine learning methods. One approach is to construct lightweight surrogate models to substitute the simulations used in inference. Alternatively, one might altogether circumnavigate the need for Bayesian sampling schemes and directly estimate the posterior distribution. We focus on stochastic simulations that track autonomous agents and present two case studies of real-world applications: tumour growths and the spread of infectious diseases. We demonstrate that good accuracy in inference can be achieved with a relatively small number of simulations, making our machine learning approaches orders of magnitude faster than classical simulation-based methods that rely on sampling the parameter space. However, we find that while some methods generally produce more robust results than others, no algorithm offers a one-size-fits-all solution when attempting to infer model parameters from observations. Instead, one must choose the inference technique with the specific real-world application in mind. The stochastic nature of the considered real-world phenomena poses an additional challenge that can become insurmountable for some approaches. Overall, we find machine learning approaches that create direct inference machines to be promising for real-world applications. We present our findings as general guidelines for modelling practitioners.

**Author summary:** Computer simulations play a vital role in modern science as they are commonly used to compare theory with observations. One can thus infer the properties of a observed system by comparing the data to the predicted behaviour in different scenarios. Each of these scenarios corresponds to a simulation with slightly different settings. However, since real-world problems are highly complex, the simulations often require extensive computational resources, making direct comparisons with data challenging, if not insurmountable. It is, therefore, necessary to resort to inference methods that mitigate this issue, but it is not clear-cut what path to choose for any specific research problem. In this paper, we provide general guidelines for how to make this choice. We do so by studying examples from oncology and epidemiology and by taking advantage of developments in machine learning. More specifically, we focus on simulations that track the behaviour of autonomous agents, such as single cells or individuals. We show that the best way forward is problem-dependent and highlight the methods that yield the most robust results across the different case studies. We demonstrate that these methods are highly promising and produce reliable results in a small fraction of the time required by classic approaches that rely on comparisons between data and individual simulations. Rather than relying on a single inference technique, we recommend employing several methods and selecting the most reliable based on predetermined criteria.

## Introduction

Mathematical and computational modelling opens up and sheds light on a wide variety of research questions in fields, ranging from hydrodynamics to oncology [1–4]. They have thus become crucial to advances in nearly all research areas and for addressing a variety of real-world problems. However, in many cases, detailed mechanistic mathematical models of nature are notoriously complex and computationally cumbersome. When faced with such models, direct statistical analyses can become insurmountable due to the associated computational cost. In this paper, we address ways to mitigate this drawback.

We take a look at a specific subset of heavy computational simulations, known as agent-based models (ABMs) [5]. Such models keep track of autonomous agents — be it, individuals, cells, or particles — that follow a set of rules prescribing their behaviour relevant to the particular phenomenon being modelled. Importantly, these rules are oftentimes stochastic and involve both interactions among the agents and between the agents and their environment. If a simulation is repeated, its output, therefore, changes in complex ways. Before one might compare the predictions of such ABMs to observations, one must gauge this stochasticity, which might consume considerable computational resources. Even without this additional complication, proper (Bayesian) inference requires a large number of simulations to ensure a robust exploration of the model’s parameter space, which could be large [6]. However, ABMs for real-world applications often require many agents or complex interactions making simulations computationally demanding. Due to the high computational cost, a direct route for simulation-based inference, such as Approximate Bayesian computation (ABC) methods [7, 8], might thus be impracticable.

There are different problem-dependent ways to deal with the computational challenges that complex inference problems pose. Some models lend themselves to statistical techniques that greatly reduce the number of required simulations [9]. In other cases, one might resort to interpolation in an existing archive of models or to parameterizations of emergent properties [10, 11]. In recent years, it has become increasingly popular to train machine learning (ML) methods to tackle the issue [12, 13]. We follow this latter approach here.

ML schemes can be employed in two different ways. First, one can create a computationally efficient surrogate model, a so-called emulator, that mimics relevant aspects of the ABM, much like the aforementioned parameterizations [14, 15]. The emulator might then enter in a Bayesian framework, such as ABC, in the same way, that the original simulations would. Secondly, one might directly use ML approaches for inference [13]. We will refer to this latter method as a direct inference machine. Whether one considers using emulators or direct inference machines, one is left with the choice of the specific ML algorithm. However, it is by no means clear-cut what the optimal choice is for a specific research question.

The recent literature takes advantage of advances in density estimation by neural networks and focuses on methods that estimate the likelihood or posterior density (or ratio) [12, 13]. In this paper, we take an approach that is less explored based on an approximation to Bayesian neural networks [16, 17]. This approach allows generating samples from the likelihood (a stochastic surrogate model) or from the posterior distribution rather than estimating their density. We compare the performance of this approach with methods based on Gaussian processes [18, 19] and a mixture density network [20, 21].

For this purpose, we consider two examples of real-world applications of ABMs: tumour development and the spread of infectious diseases. ABMs are used extensively in both fields [22–27]. The employed ABMs are complex enough to resemble real-world problems going beyond toy models. In both cases, we create synthetic data (also called mock observations) using the ABMs and infer the underlying model parameters using a broad range of inference algorithms. The mock data in our examples is considered experimentally accessible. This is a deliberate choice since real-world applications must always keep data availability in mind. Observations play first fiddle.

In the following, we present several implementations of emulation-based inference and direct inference machines. We apply these inference methods to the aforementioned ABMs and critically assess the results from each method. While we focus on these two examples, cancer and infectious diseases, we aim to provide general guidance for choosing which algorithm to use when dealing with computationally heavy simulations.

## Results

We use two main approaches to simulation-based inference: emulation-based approaches and direct inference machines. We employ three different emulators: a deep neural network (NN), a mixture density network (MDN), and Gaussian processes (GP). These methods were chosen as they can mimic the stochastic nature of the ABMs. The predictions of the emulators are directly compared to the mock observations, i.e. the synthetic data. We then infer the underlying model parameters (**Θ**) using rejection ABC and Markov Chain Monte Carlo (MCMC). Finally, we compare these ML predictions with the ground truth (**Θ**_o_). For the ABC, we employ a single distance measure that compares individual predictions to the mock observations, while we take two distinct approaches for the MCMC: The likelihood either relies on individual predictions (case a) or global properties of the predicted distributions (case b). With regards to the direct inference machines, we again employ a NN and GP and compare the predictions to the ground truth. For each method, we vary the size of the training set to evaluate its impact on the results as described in the following. Further details are given in the methods section.

### Comparing emulators

When comparing the predictions of the emulators to our synthetic data, we consider the absolute error in the mean, the relative error in the median, the ratio between the predicted and true width of the distribution in terms of the standard deviation, and the Wasserstein distance between predicted and true distribution (see methods section for details).

Figures 1 and 2 summarize the comparisons between the emulation-based predictions and the distributions obtained from simulations for the ABMs describing tumour growth and disease spread, respectively. Our ABM describing brain cancer is a so-called cellular automata (CA) model (cf. methods section). Our epidemiological model is a so-called SIRDS model (cf. methods section). In the following, we refer to the ABMs using this terminology.

**Fig 1.**
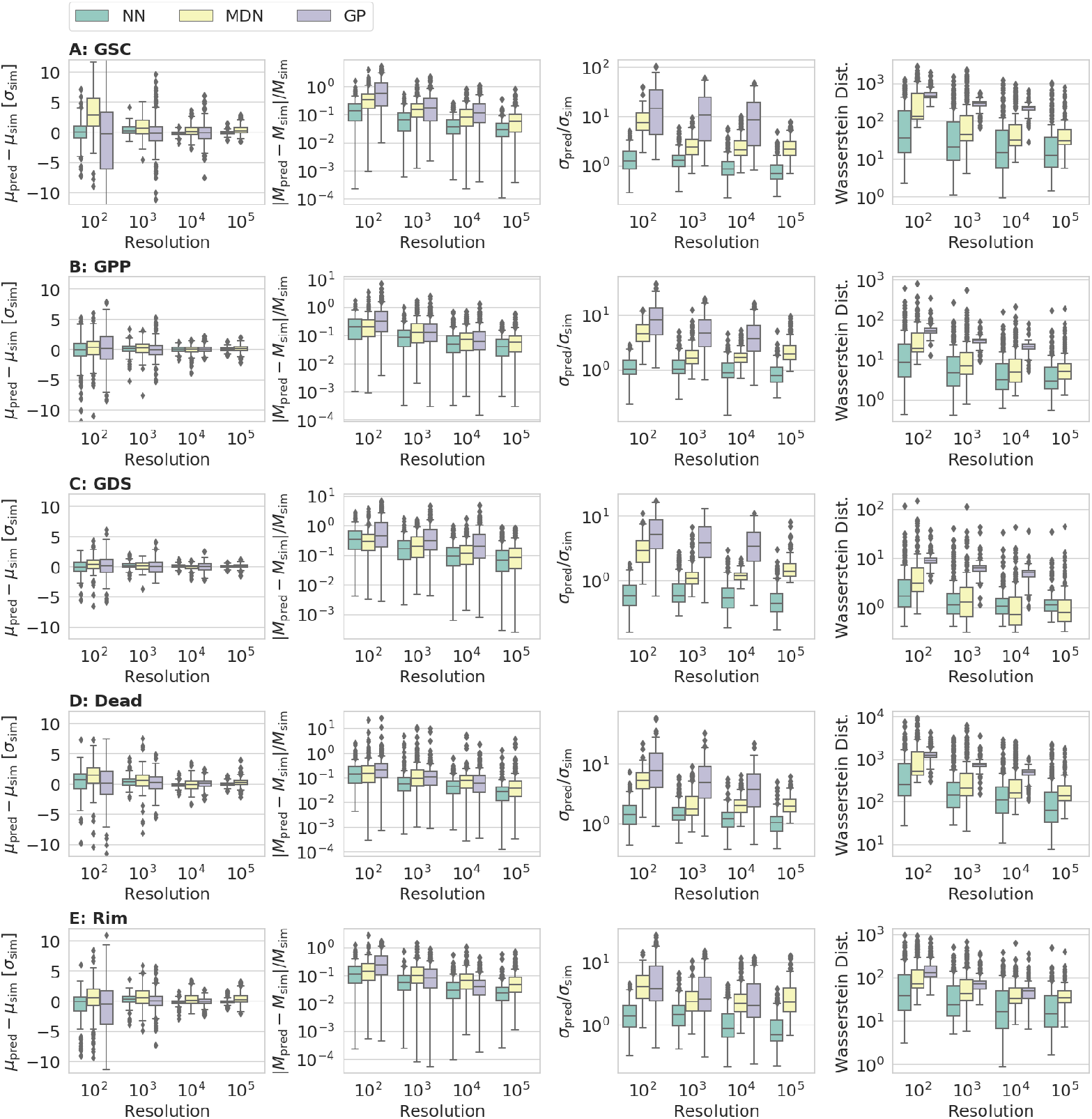
Comparison of the performance of the emulators as a function of the size of the training set (resolution) for the agent-based cellular automata (CA) model of brain tumours. The plot shows the results for three emulators: A neural network (NN), a mixture density network (MDN), and a Gaussian process (GP). The subscripts ‘pred’ and ‘sim’ refer to the predictions by the emulators and the ground truth obtained from simulations, i.e. the mock observations, respectively. From left to right, the columns contain the error in the predicted mean (*µ*_pred_), the relative absolute error in the predicted median (*M*_pred_), the ratio between the predicted width of the marginals in terms of the standard deviation (*σ*_pred_) and true width, and the Wasserstein distance (cf. Section *Metrics*). Rows A-E contain the number of glioblastoma stem-like cells (GSC), the number of cells in the propagating progeny (GPP), the number of cells in the differentiating subpopulation (GDS), the number of dead tumour cells, and the number of cells in the proliferating rim, respectively.

**Fig 2.**
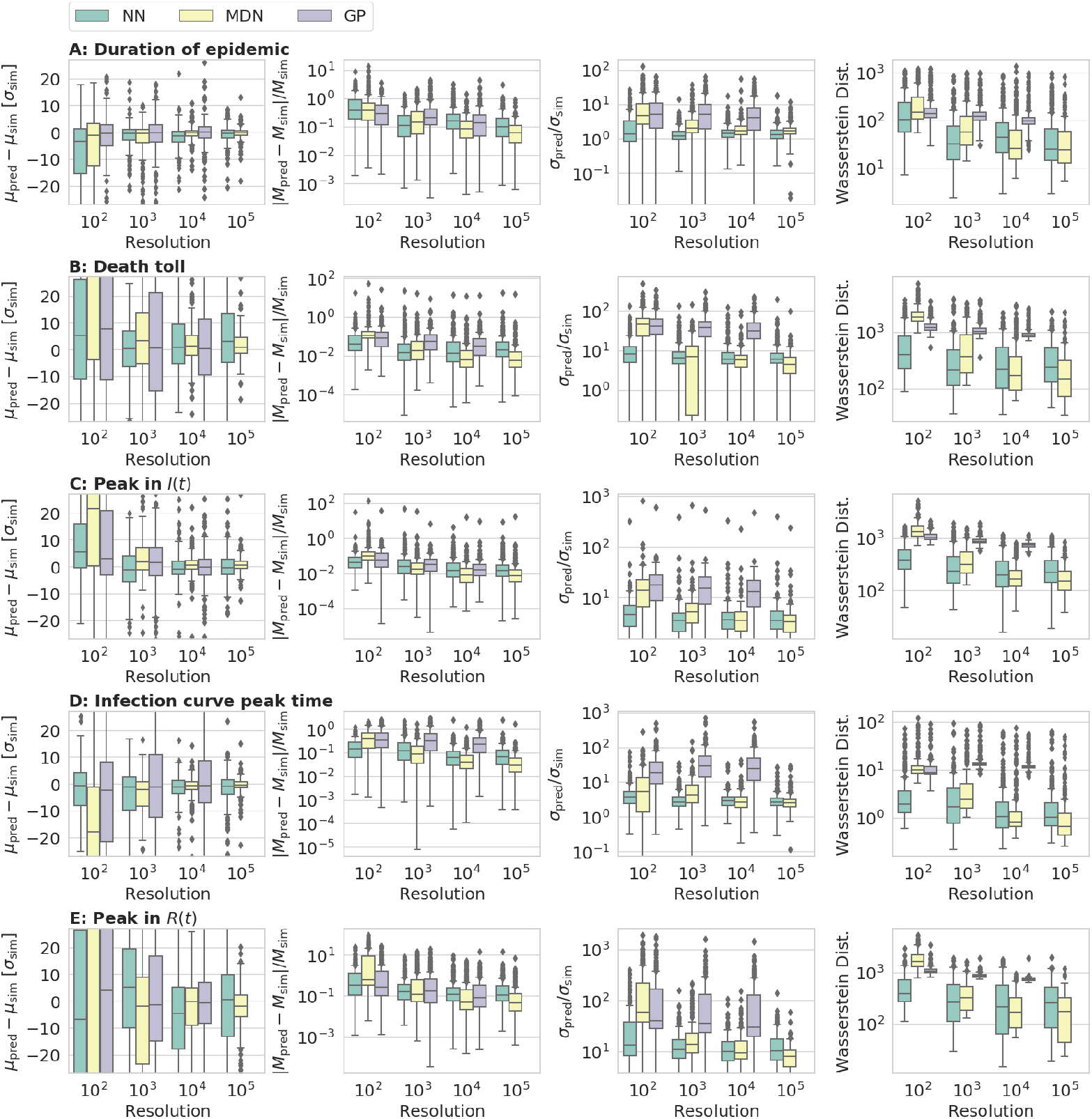
Counterpart to Fig 1 for our epidemiological SIRDS model. Rows A-E contain the total duration of the epidemic, the total death toll, the highest number of infected individuals during the outbreak, the time at which the peak in infections occurs, and the highest number of recovered agents during the outbreak, respectively.

As regards the cancer CA model for tumour growth, the NN outperforms the other ML techniques across nearly all measures, sizes of the training set, and variables. The NN closely recovers the correct mean, median, and width of the marginal distributions obtained from simulations and generally yields the lowest Wasserstein distance. In contrast, the MDN is generally biased, overestimating the true mean, while the GP results in distributions that are much too broad.

For our epidemiological SIRDS model, on the other hand, the MDN outperforms the NN for sufficiently large training sets. The GP still yields the worst performance across all measures.

As can be seen from both figures, the accuracy and precision of the different ML approaches generally improve with increasing size of the training set. However, the GP cannot cope with the kernel matrix for the largest training set. This is not to say that this is a general flaw of Gaussian processes [28] but rather goes to show that the suitability of any ML implementation or approach is problem-dependent.

### Comparing inference techniques

When comparing the inference results to the ground truth, we include four metrics: We compare the mean of the distribution to the true parameters and compute both the relative and absolute error. Moreover, we consider the standard deviation of the posterior and the probability density (*q*) attributed to the true value (see further details on metrics used in the method section).

As mentioned above, the NN and GP result in the best and worst performance among the explored emulators for the cancer CA model, respectively. To see what impact this performance gap has on the inference, we sampled the parameter space using both emulators in tandem with both ABC and MCMC. Due to the associated computational cost, we do not repeat this analysis for different sizes of the training set. Instead, we settle for a resolution of 10^4^ simulations in the training set, as the NN performs well at this resolution. Any additional gain in performance at 10^5^ simulations does not outweigh the substantially increased computational expense. For our epidemiological SIRDS model, we limit ourselves to the NN as regards the emulation-based inference.

For the cancer CA model, we find that the higher accuracy and precision of the NN emulator is reflected in the performance metrics for the inference results. We illustrate this result in Fig 3. Whether we use the ABC or MCMC algorithm for inference, the NN leads to a narrower distribution, whose mean more closely recover the ground truth than the GP does.

**Fig 3.**
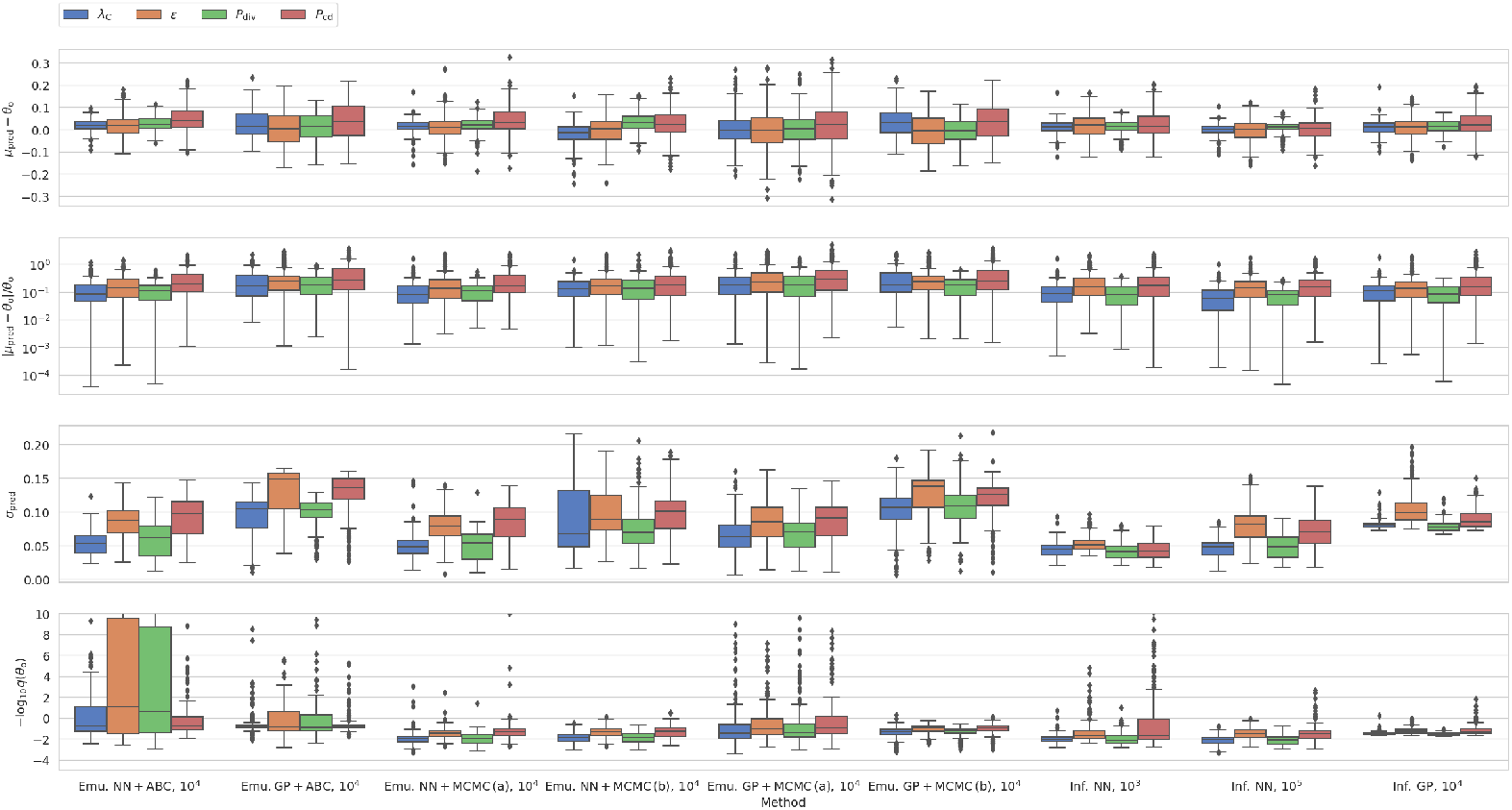
Performance across different inference schemes for our cancer CA model. The first row contains the residuals between the mean of the marginal distributions and the true parameter values (Θ_o_) for each method. The second row shows the corresponding relative error in the prediction. The third row summarizes the standard deviation of the marginals, while the fourth row shows the negative logarithm of the probability density (*q*) attributed to the true parameter value. The plot includes the results from the emulation-based approaches (emu.) as well as our direct inference machines (inf.). We include two machine learning approaches: A neural network (NN) and Gaussian processes (GP). In connection with the emulators, we distinguish between results obtained when using rejection ABC and MCMC. For each approach, the label specifies the size of the training set: For the emulators, we consistently used 10^4^ simulations.

To put the results in Fig 3 into perspective, consider the case where we simply use the mean of the posterior distribution as our parameter estimate. This approach would result in unbiased residuals for the mean with 25th and 75th percentiles at −0.1 and 0.1, respectively. All methods thus perform better than a random assignment of parameter estimates.

Irrespective of the ML approach, the MCMC (cases a and b) and ABC lead to similar values of the four measures, including −log *q*(*θ*_*o*_). However, due to the stochastic nature of the emulator, we note that the median autocorrelation lengths across all four input parameters are 383-412 and 1889-1953 samples for NN and GP MCMC runs, respectively, when individual predictions by the emulator are used in the likelihood (case a). We summarize these numbers in Table 1. The GP emulator thus results in a lower number of effective samples. Indeed, for the GP, the resulting contour plots are very noisy, often consisting of disconnected peaks. This notion calls the robustness of the MCMC results based on the GP into question. We attribute this to the fact that the GP emulator overestimates the standard deviation for the output parameters of the ABM. This increased stochasticity hinders convergence.

**Table 1.** Median and mean autocorrelation length for different MCMC setups. For each of the two ABMs (our cancer CA and epidemiological SIRDS model), we use two ML emulation algorithms (NN and GP). We distinguish between two different likelihoods (case a and b, cf. the methods section). The first column specifies the different setups by first naming the ABM followed by the emulator and the likelihood.

| MCMC | Mean | Median |
| --- | --- | --- |
| CA-NN (a) | 565-599 | 383-412 |
| CA-NN (b) | 377-402 | 163-174 |
| CA-GP (a) | 1835-1875 | 1889-1953 |
| CA-GP (b) | 423-456 | 311-346 |
| SIRDS-NN (a) | 2084-2120 | 2118-2179 |
| SIRDS-NN (b) | 325-393 | 194-266 |

As regards the epidemiological SIRDS ABM, the case a of the MCMC algorithm suffers from the same issue, even when dealing with emulators that closely recover the marginal distributions obtained from simulations. For the NN MCMC runs, the mean and median estimated autocorrelation lengths across the different parameters lie between 2084 and 2179, and the associated contour plots are extremely noisy when individual predictions by the emulator are used in the likelihood (case a). Due to this behaviour, we have omitted case a of the MCMC results in Fig 4 that summarizes the performance of the different inference techniques for the epidemiological SIRDS ABM.

**Fig 4.**
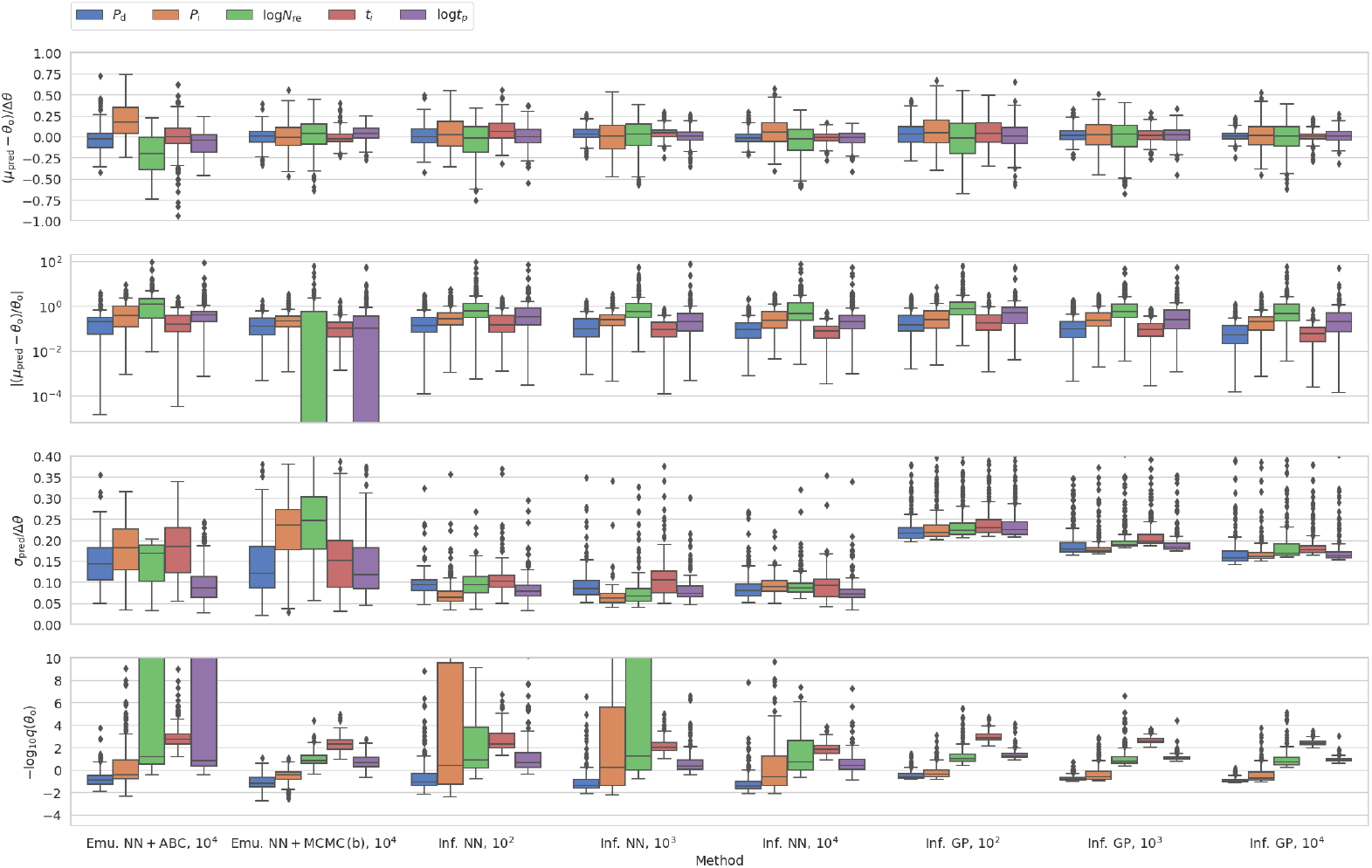
Counterpart to Fig 3 for our epidemiological SIRDS model.

Rejection ABC, on the other hand, does not suffer from the same problem for either of the two ABMs. Even though rejection ABC leads to high values of −log *q*(*θ*_*o*_) in some cases, this technique is able to reliably reconstruct the mean and median. Moreover, in contrast to the MCMC that leads to spurious contour plots, rejection ABC does not yield equally noisy posteriors even though the distance measure draws on individual predictions by the emulator. To improve the contour plots of the MCMC, we have to resort to using global parameters of the predicted distributions rather than individual samples in the likelihood (case b). When doing so, the median autocorrelation length lies in the intervals 163-174 and 194-266 for the cancer CA and epidemiological SIRDS models, respectively (cf. Table 1).

Figures 3 and 4 also includes the predictions by the NN and GP direct inference machines based on different sizes of the training set. Two features stand out: First, the direct inference machines perform as well or better than the emulation-based approaches, even with a low number of simulations in the training set. Indeed, some measures do not visibly improve even when the model is confronted with significantly more training data. Secondly, the dissonance in −log *q*(*θ*_*o*_) between the NN and GP is more prominent for the epidemiological SIRDS ABM than in the case of the cancer CA model.

As regards both ABMs, it bears mentioning that the values obtained for the four metrics are partly correlated. However, the correlations are method- and measure-dependent. They might thus be erased when comparing samples across different methods and measures, reflecting e.g. differences in the performance and the normalization factors.

Finally, as can be seen from the figures, all inference techniques, in broad strokes, struggle with the same model parameters, while they are all reasonably good at establishing certain other model parameters. This behaviour is hardly surprising. It merely reflects the non-identifiability of certain parameters given the information that is available through the summary statistics. However, exactly because this behaviour is to be expected, it provides an additional sanity check, suggesting consistency of the different approaches. While one might overcome this issue by choosing different summary statistics, we note that this might not be straightforward in real-world applications — e.g. due to limitations imposed by the data.

### Performance analysis

To quantify the motivation for using the different inference techniques, we summarize the CPU time requirements for each method in Fig 5. Of course, the computation time is greatly model-dependent. Our intention is thus merely to underline a few key features.

**Fig 5.**
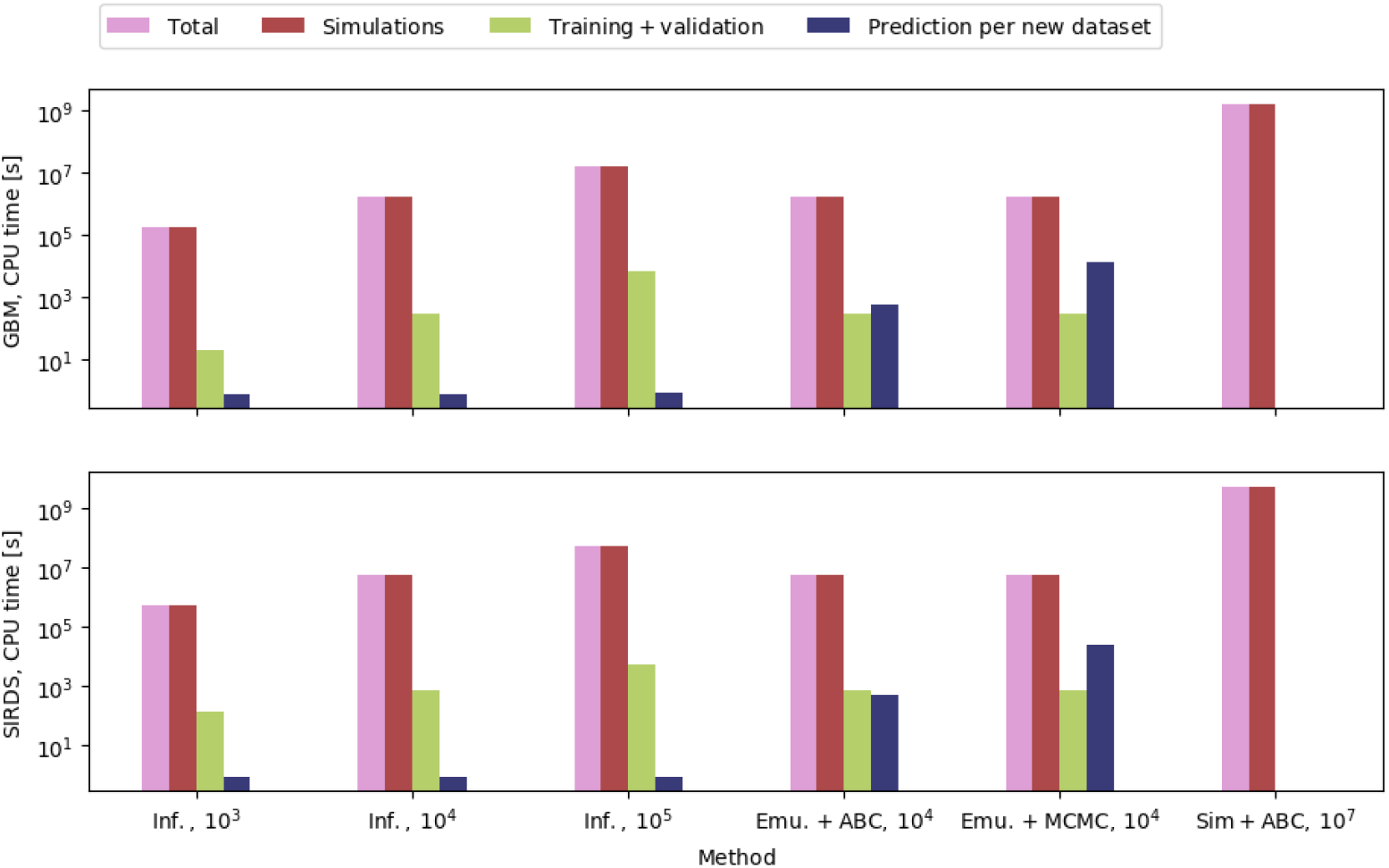
Required CPU time across different methods for our cancer CA model (upper panel) and the epidemiological SIRDS model (lower panel). The light magenta bars show the total time required for each inference technique. The remaining three bars specify the time required for the simulations involved (red), the time required for training and validation (yellow), and the time that is needed to infer the parameters for a single set of observations (blue). The number of required simulations is specified in the labels. We include the NN direct inference machine based on three different sizes of the training set. Moreover, we include the NN emulator in tandem with the rejection ABC and MCMC algorithms. The ABC accepts the best 10^4^ among 10^7^ randomly drawn samples. The ensemble MCMC uses 8 walkers, drawing 20,000 samples for each walker. For the MCMC, we average over the CPU times for cases a and b. For comparison, we include an estimate on the time that would be required for running the ABC with 10^7^ samples using the actual simulations. Note that the ordinate is logarithmic. Note also that the time needed to run any given simulation depends on the number of agents and hence the model parameters.

As can be seen from the figure, the computation of the actual simulations by far consumes the most resources (the total time in every case is almost equal to the simulation time). Training the direct inference machines or the emulators takes orders of magnitudes less time. Due to the nature of the ABMs, any method that reduces the required number of simulations thus has an edge. This also illustrates why we do not favour inference methods that rely on the actual simulations, especially if new simulations would be required for every new set of observations — MCMC algorithms are a good example of a method that require new simulations for each set of observations, while rejection ABC and other grid-based methods [29] do not suffer from this issue. Indeed, even with simplistic ABMs, such a task might become computationally insuperable. In contrast, when dealing with the direct inference machines or the emulators, new simulations are only required during the training phase.

For our models, the training of the emulators and direct inference machines take the same time, which does not always have to be the case. However, the direct inference machines are more effective when it comes to obtaining estimates for the underlying model parameters. Moreover, since the simulations take the most time, investing in training more complex ML emulators or direct inference machines will not be overall more computationally expensive and may pay off in the accuracy of the results.

## Discussion

In this paper, we investigate how to efficiently perform inference when relying on computationally expensive stochastic agent-based models (ABMs). For this purpose, we compute synthetic data and infer the underlying model parameters using a variety of techniques. These methods fall into two main categories: emulators and direct inference machines. We find that the performance of the emulators and direct inference machines are not consistent across the two ABMs but rather depend on the real-world application in question as well as on the size of the training set and thus on the available computational resources. This leads us to our first guideline:

- **There is no one-size-fits-all solution for simulation-based inference**. The performance of any given algorithm is problem-dependent. Since there are *a priori* no clear-cut preferences, we recommend confronting data with several techniques and selecting the best method based on suitable metrics.

We do hence not present an absolute ranking of the methods per se but rather raise caution about solely relying on a specific method or implementation when facing a new research problem. We do, however, note that the GP performed worse than the NN and MDN methods. So, one may need to be careful when relying on Gaussian processes for emulation [30, 31]. This being said, other implementations of Gaussian processes might be able to address the shortcomings of the implementation used here [32, 33].

The notion that different methods may work better for different problems is consistent with the findings by Lueckmann et al. [13], who study a range of benchmark problems to provide guidelines for so-called sequential inference techniques. In a nutshell, sequential techniques explore the parameter space in an iterative fashion. Rather than relying on a static grid for training, they hand-pick those points in the parameter space that closely match the specific observation whose properties we aim to infer. Sequential techniques are hereby able to greatly reduce the number of required simulations by orders of magnitude. Implementations for both emulation and direct inference exist [34, 35]. The catch is that new simulations are required for every new set of observations. With 250 test cases for each ABM in our analysis, we can thus not reap the benefits of these algorithms as a substitute for our inference machines. However, sequential techniques have proven very powerful in fields, such as cosmology [36]. After all, there is only *one* universe.

When comparing our results to those by Lueckmann et al., it is also worth noting that we use an alternative application of neural networks for emulation and direct inference in this paper: Rather than estimating the likelihood or posterior densities, we use a neural network with dropout that provides an approximate Bayesian formulation and mimics a Gaussian process [16, 16, 17, 37, 38]. Like the methods discussed by Lueckmann et al., we find this alternative approach to be very effective in producing good emulators and direct inference machines with a relatively small number of simulations sampled from the parameter space in our examples.

Yet another other distinct aspect of our study is that we keep the limitations of the real-world ABM applications in mind: When dealing with real-world data, one must often resort to simplifying assumptions regarding the true distribution of the data, due to the nature of the experiments. Not only does this dictate the type of data that we can gather but it also imposes constraints on the summary statistics and the metrics that we can draw on (cf. the methods section). Our paper outlines how one might go about selecting different methods and performance measures based on the research problem using ABMs as a case study.

We do by no means claim that our list of emulators and direct inference machines is exhaustive. Other viable approaches exist with scope for future developments. For example, recent work on generating differential equations from stochastic ABMs [39] might be extended to stochastic settings to generate new emulators. However, it is beyond the scope of the present paper to explore all possible paths. Rather, we intend to present an illustrative selection. We also note that not all ML methods can readily address the stochastic nature of the data. At least not in their standard form. Acquiring error bars might, oftentimes, require elaborate extensions (see e.g. the paper by Meinshausen et al. [40] for a discussion on regression random forests [41]).

For our ABMs, we find that the direct inference machines provide accurate and precise results even with a very limited training set. Since computation time is one of the most prominent obstacles when dealing with ABMs for real-world applications, this feature favours direct inference algorithms. Thus,

- we generally recommend to **look for techniques that minimize the number of simulations required** such as direct inference machines.

This is not to say that direct inference machines will be the optimal choice for all ABMs. The present success of the direct inference machines might reflect aspects, such as the complexity of the true posterior distributions or the adequacy of the distance measure and likelihood used by the statistical inference technique, i.e. rejection ABC and MCMC. However, we note that the presented direct inference machines (as well as the emulation-based ABC or MCMC) can produce multi-modal posteriors.

As regards the statistical inference techniques, we find that the MCMC does not always yield robust results. We attribute this to the stochastic nature of the ABMs. While the rejection ABC consistently yields worse results than the direct inference machines in terms of the employed metrics, it does not suffer from this shortcoming.

- Due to the stochastic nature of the agent-based models, **some statistical inference techniques do not yield robust constraints for emulation-based approaches**.

The statistical algorithms explored in this paper, i.e. rejection ABC and MCMC, are by no means exhaustive. Other viable alternatives, such as sequential Monte Carlo (SMC) [42], exist. The aforementioned sequential techniques discussed by Lueckmann et al. offer another alternative when combined with the emulators. However, we also note that the stochastic nature and intractable likelihood of the explored systems render some approaches impractical. For instance, while Hamiltonian Monte Carlo (HMC) [43] is generally a demonstrably powerful technique [44], it is not suited for our purposes as we cannot reliably compute derivatives in the explored parameter space.

By addressing and comparing multiple widely-used algorithms, we hope that the present paper assists researchers in selecting appropriate inference techniques for real-world applications. While we have focused on ABMs here as one class of real-world models, we believe our results to be relevant to other stochastic complex systems. State-of-the-art numerical models are often computationally heavy, and we hope that our guidelines and recommendations help modellers to overcome this stumbling block. To facilitate this goal, we have made our code available on https://github.com/ASoelvsten/ABM.

## Methods and Models

### Agent-based models

In this paper, we use two agent-based models (ABMs) describing two distinct real-world problems: Our first model deals with a malignant type of brain cancer called glioblastoma multiforme. The second model describes the spread of infectious diseases in a population. The appendices *Brain tumour, CA model* and *Epidemic, SIRDS model* provide detailed accounts of both ABMs. Since these models are solely to be understood as examples that we have used to benchmark different inference methods, we limit ourselves to remark on a few key aspects of the models here. In both models, the agents live in a two-dimensional plane, and the dynamics of the system is governed by a set of stochastic rules that dictate the behaviour of the individual agents.

### Brain tumour, CA model

Our agent-based brain cancer model is a so-called cellular automata (CA) model, in which each agent represents a tumour cell. This ABM has four input parameters. Our first two parameters, *P*_div_ and *P*_cd_, are probabilities that are associated with the rules for cell division and spontaneous cell death. The third parameter, *λ*_C_, determines the nutrient consumption of the cells and entails the possibility of cell death due to nutrient deprivation. This parameter is a rate given in units of the nutrient consumption per cell per time step.

To explain the fourth parameter, we need to mention that brain tumours are composed of three main different cell types [45]. In the following, we refer to these as glioblastoma stem-like cells (GSC), glioblastoma cells in the propagating progeny (GPP), and glioblastoma cells in the differentiating subpopulation (GDS), respectively. The fourth parameter, *ϵ*, thus denotes a probability that is associated with the differentiation of stem-like cells during cell division. All four parameters are varied on a linear scale from 0.01 to 0.50 (see the appendix *Parameters and summary statistics*).

In this paper, we perform inference based on synthetic data, i.e. mock observations. To do so, we need to consider what data would be available in a real-world scenario. In brain cancer research, data often comprise detailed snapshots of tumour growth. To reflect this circumstance, we assume that we know the time frame during which the tumour has evolved. For the purpose of inference, we hence collect data from all simulations at a fixed snapshot (*t* = 100). To compare the model predictions to the synthetic observations, we focus on the emergent macroscopic properties of the ABM. As our summary statistics, we use five numbers describing the system as a whole: the number of GSCs, GPPs, GDSs, and dead tumour cells as well as the total number of alive cells in the so-called proliferating rim (cf. the appendix *Brain tumour, CA model*).

### Epidemic, SIRDS model

In our model for the spread of infectious diseases, the agents are people. These individuals fall into four different groups as signified in the acronym of the model: SIRDS. The first group comprise the people who are susceptible (*S*) to the disease. The second group includes people who are infected (*I*). In the third group, we find those people who are immune after having recovered (*R*) from the disease. The last group includes those who have died (*D*) from the disease. As the infection spreads through the population, individuals move between groups.

Our SIRDS model has five input variables. The first parameter, *P*_i_, denotes the probability of contracting the disease upon contact with an infected individual. The second parameter, *P*_d_, denotes the probability of dying from the disease. The agents have fixed social relations (at home, at school, or at work). In addition, they have *N*_re_ random encounters. The logarithm of *N*_re_ constitutes our third parameter. As our fourth parameter, we choose the number of days (*t*_i_) that an infected individual is contagious. Finally, our fifth parameter is the logarithm of the length of time, log *t*_p_, during which a recovered individual is protected from getting the disease. After this time span, the previously recovered individual becomes susceptible (*S*) again.

Note that the parameters of our SIRDS model are different from one another in terms of the explored values, i.e. the priors. This stands in stark contrast to the parameters of the cancer CA model. Also, the models are quite distinct in terms of mechanistic details (See the appendices *Brain tumour, CA model* and *Epidemic, SIRDS model* for details). Thus, by construction, the cancer CA and SIRDS models complement each other.

As regards the summary statistics, we again resort to the emergent properties of the system. Rather than relying on a single detailed snapshot, epidemiologists often have comprehensive knowledge of *D*(*t*), *I*(*t*), and *R*(*t*). Here, we boil these time series down to five numbers: the total duration of the epidemic, the total death toll, the highest number of infected individuals during the outbreak, the time at which the peak in infections occurs, and the highest number of recovered agents during the outbreak (cf. the appendix *Parameters and summary statistics*).

We stress that both our cancer CA and our epidemiological SIRDS simulations are rather simplistic. However, they are sufficiently realistic to capture the main aspects of the real-world applications, including the stochastic nature of the modelled events. This feature is the most essential for our purposes. Rather than providing detailed models for specific biological systems, we investigate how to infer properties from observations based on such simulations. By keeping the models simple, we lower the computational cost making the analysis more practical and allowing us to readily explore different aspects.

### Model grid

To perform inference, we construct a set of grids in the input parameter spaces with a limited number of simulations. We then use these to train the machine learning (ML) techniques discussed in the sections *Emulators* and *Direct inference* below.

We impose uniform priors on all parameters. Our parameter spaces are thus four and five-dimensional hypercubes for the cancer CA and epidemiological SIRDS model, respectively. We could cover the parameter spaces by sampling points in regular intervals. However, this approach is rather inefficient. Alternatively, we could generate points randomly. However, this approach leads to unintended clustering. We, therefore, follow the suggestion by Bellinger et al. [46] and generate points based on a Sobol sequence [47]. Doing so, we sample the hyperspace uniformly, while minimizing redundant information as discussed by Bellinger et al. [46].

Table 2 lists the lower and upper boundaries of our priors, i.e. the boundaries of the sampled parameter space.

**Table 2.**
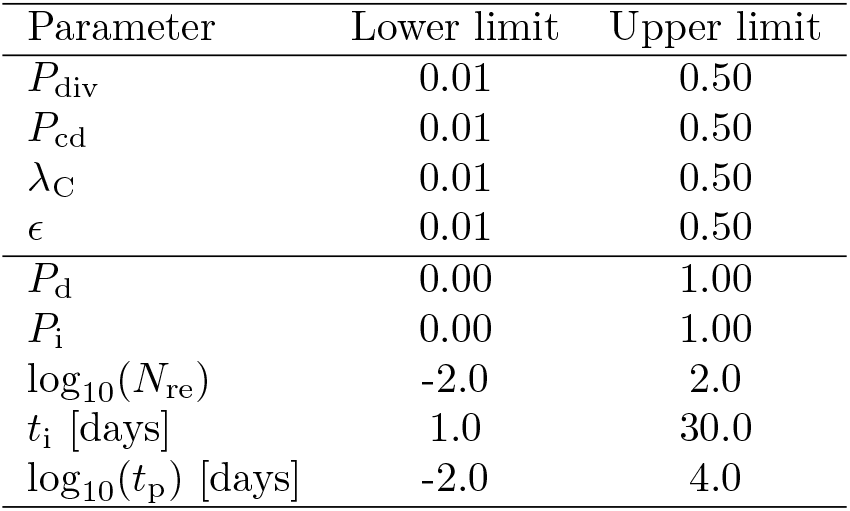
Upper and lower boundaries for the model parameters (Θ) in the constructed grids.

For both ABMs, we construct four grids with different resolutions. These four grids contain 10^2^, 10^3^, 10^4^, and 10^5^ simulations, respectively. Each simulation has a unique parameter set, i.e. the grids do not contain repetitions. Rather, the ML algorithms must infer the stochasticity of the simulations based on the variation between neighbouring simulations across the grid.

Since we cover a relatively broad region of each parameter space, we run into scenarios, where, say, all tumour cells die before *t* = 100, or where the disease is not passed on. As discussed in the section *Mock observations*, these scenarios are of no interest. They are, therefore, effectively noise for the ML algorithms. By excluding them from the training sets, we boost the performance of the various inference techniques. More specifically, in connection with the cancer CA model of brain tumours, we remove all simulations, for which the number of GSCs or GPPs is zero. As regards our epidemiological SIRDS model, we remove all cases, in which the disease did not spread beyond the originally infected individuals or the disease was not eradicated after 1825 days (5 years). The latter exclusion criterion stems from the simulation set-up (cf. the appendix *Epidemic, SIRDS model*). For both ABMs, these criteria reduce the original grid sizes by roughly 10 per cent.

### Mock observations

We construct a synthetic dataset to discuss the performance of the different inference techniques. For each ABM, we generate 250 new input parameter sets and repeat each simulation 250 times to sample the resulting distributions of the output variables. These distributions can be compared to the predictions of the emulators discussed in the section *Emulators*, while the original input parameters (**Θ**_*o*_) of the simulations can be compared to the posterior distributions obtained by inference. We present the metrics for these comparisons in the section *Metrics* below.

We pick the model parameter values for our mock observations using tighter priors than those presented in Table 2. The benefits of this approach are twofold. First, we avoid drawing simulations at the edges of the regions covered by the grids. Secondly, by choosing different priors, we select points that differ from those that enter the training data although we use a Sobol sequence to sample the parameter space.

In the section *Results*, we infer the properties of our mock observations using various techniques and compare the results with the ground truth. This comparison allows us to assess the performance of the different inference algorithms. However, due to the distance measure and likelihoods associated with the employed Bayesian inference techniques, i.e. ABC and MCMC, not all points in the mock observations might be equally suitable for our purposes (cf. the sections *Rejection ABC* and *MCMC*). Especially when one or more of the output variables consistently take on a fixed value, the defined distance measure and likelihood functions run into trouble since we end up dividing by zero. One such scenario would be the case where the parameter values ensure that all tumour cells consistently die off before *t* = 100. Another example is the case where the disease is never passed on. To mitigate this problem, one might discard all mock observations that produce delta functions as the marginals for one or more output variables of the ABMs — in the real world, such scenarios would not be studied, anyway. Here, we apply a more conservative criterion, discarding all mock observations, for which the width of the 68 per cent confidence interval is zero. With this selection criterion, we are left with 219 and 227 points in the parameter space for the cancer CA and epidemiological SIRDS models, respectively.

### Emulators

The purpose of an emulator is to mimic the output of the ABM at low computational cost when given a set of ABM input parameters, often called features. Like the original ABM, the emulator is thus a function *f* (**Θ**) that maps from the model parameter space (**Θ**) to the output space of the ABM. The emulator can then be used as a surrogate model to sample the posterior probability distributions of the model parameters using likelihood-free inference (LFI) or other Bayesian inference techniques (cf. the sections *Rejection ABC* and *MCMC*).

Since the ABMs in this paper are driven by stochastic events, several simulations with the *same* input parameters will produce a distribution for each output variable of the ABM rather than a unique deterministic result. The emulator must capture this behaviour. Based on this notion, we follow three different approaches: We emulate the simulation output using a deep neural network (NN), a mixture density network (MDN), and Gaussian processes (GP).

We train and validate the emulators based on precomputed grids of models (cf. the section *Model grid*). When the ML methods rely on neural networks (NN and MDN), 60 per cent of the simulations are used for training, while the remaining 40 per cent are used to validate and test the methods before they encounter the mock observations (cf. the sections *Mock observations* and *Results*). For the Gaussian processes, 20 per cent are set aside for a testing phase, while the remaining 80 per cent of the simulations are used for training — when dealing with neural networks, the validation set is e.g. needed for invoking early stopping during training. We also note that the different ML approaches do not see the raw mock observations: Since we are working with outputs on different scales, the mock observations are standardized before handing it over to the ML schemes.

### Neural networks

As our first approach, we construct a broad deep neural network (NN) with 3 hidden layers that contain 100 neurons each. To avoid overfitting, we include early stopping as well as dropout during the training phase [48]. The latter property implies that we randomly select 20 per cent of the neurons in each layer and temporarily mask these neurons during each pass through the NN [48]. To recover the stochastic nature of the ABMs, we maintain dropout during the inference phase and use the loss function proposed by Gal & Ghahramani [16, 37, 38]. These temporal changes in the NN’s architecture mean that the predictions of the NN become samples from a distribution rather than being deterministic. The resulting distribution encodes both epistemic and aleatoric uncertainties, i.e. those stemming from the limited knowledge of the NN and those stemming from the stochasticity of the ABMs. NNs that employ dropout have been shown to approximate Gaussian processes [16] and can be seen as an approximation of Bayesian neural networks [17]. Our implementation builds on TensorFlow [49].

### Mixture density networks

As our second approach, we employ a mixture density network (MDN) that fits a mixture of multivariate normal distributions to the output of the ABMs [20, 21]. Within the framework of MDN, it is straightforward to mimic the stochasticity of the underlying ABM: During inference, we simply sample from the constructed mixture model.

The fit is accomplished by training a neural network to predict the means and full covariance matrices for each of these normal distributions. We use three components for each mixture model. For a five-dimensional space of output variables from the ABM, the neural network is thus optimized to predict 63 variables: 15 means, 15 diagonal and 30 unique off-diagonal elements of the covariance matrix, and 3 component weights. To ensure that the covariance matrices are positive semi-definite, we use exponential and linear activation functions for the diagonal and off-diagonal elements, respectively [36]. To guarantee that the component weights are positive and sum to unity, they are meanwhile passed through a softmax activation function. Regarding the loss function, we maximize the log-likelihood that the mixture model attributes to the training data from the grid discussed in the section *Model grid*. As in the section *Neural networks*, we employ early stopping and dropout. Our implementation builds on TensorFlow [49].

### Gaussian Processes

Finally, as our third approach, we describe the output of the ABM as a Gaussian process (GP) [18, 50]. For a comprehensive overview of GPs, we refer the reader to the work by Rasmussen et al. [19].

We use a linear combination of a radial basis function (RBF) kernel and a white noise kernel to specify the covariance of the prior function. We do so to properly account for the spread in the underlying output from the ABM [51]. Just as in the case of the MDN, the stochasticity of the ABM can be emulated during inference by sampling from the obtained GP.

Here, we set the mean squared error in the prediction to be the loss function, which amounts to the simplifying assumption that the output variables of the ABM are uncorrelated. We use the python implementation of Gaussian process regression from scikit-learn [52].

We note that the employed implementation produces a uni-modal distribution for a unique set of input parameter values. The same holds true for the NN. In contrast, the MDN is able to produce multi-modal outputs when given a single input vector.

### Rejection ABC

Having constructed an emulator, we can resort to standard sampling techniques to map the posterior probability distributions of the model parameters (**Θ**) of the ABM given a set of observations (***x***). In our case, these observations constitute the mock observations created by the ABM as discussed in the section *Mock observations*.

One option is to draw on the class of Approximate Bayesian Computation (ABC) methods [53–55]. The simplest among these algorithms is rejection ABC. Rejection ABC randomly draws samples from the prior probability distributions of the model parameters. The algorithm then compares the simulation output, i.e. the predictions of the emulator, with the observations using a suitable distance measure. Samples that lie within a distance *δ* are kept, while the rest are discarded. In practice, *δ* is chosen in such a way as to keep a certain fraction of the samples. The distribution of these samples is then taken as a proxy for the true posterior.

In this paper, we use the implementation by Lintusaari et al. [56] (elfi). As regards the prior, we use uniform priors corresponding to the region covered by the grids discussed in the section *Model grid*. As regards *δ*, we keep the best 10^4^ of 10^7^ samples for each combination of parameters for our mock observations.

The last thing that we need to specify for the ABC is the distance measure. To motivate our choice, we need to consider the nature of the real-world applications that our ABMs strive to model. At best, a targeted laboratory study of glioblastoma or disease transmission may involve a handful of animals, such as mice or ferrets [57, 58]. It can, therefore, be notoriously complicated to assess the stochasticity that underlies the data. The same holds true when dealing with patient-specific data or data from the outbreak of an epidemic [59].

For instance, consider an experiment, in which we implant human tumour cells in a single mouse to study tumour growth. Based on this time series alone, we cannot hope to rigorously constrain predictions for the outcome when repeating this experiment. Even given a handful of such experiments, we might be hard-pressed to go beyond simplifying assumptions, such as the notion that the observations will be normally distributed. Indeed, one might often not have enough information to go beyond measures, such as the median, the mean, the standard deviation, the standard deviation of the mean, the mean absolute error and/or the median absolute error. With this in mind, we settle for a distance measure (*d*) in the form

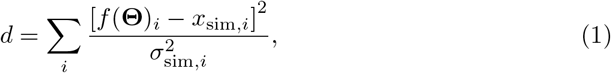

where the sum runs over all five observed output variables of the ABM, *x*_sim,*i*_ is the median of the marginal distribution for the *i*th summary statistics in the mock observations, and *σ*_sim,*i*_ denotes the standard deviation of the synthetic data. The distance is computed for each prediction made by the emulator denoted by *f* (**Θ**).

We note that *x*_sim,*i*_ and *σ*_sim,*i*_ do not fully capture the distribution of the mock observations. In other words, by employing Equation (1), we assert the notion that real-world data do not reveal the same level of detail regarding the stochasticity of the studied events as our mock observations do. This being said, we also note that the marginal distributions obtained from the mock observations are, for the most part, uni-modal and relatively symmetric distributions. The simplifying assumptions that enter Equation (1) thus still give a reasonable depiction of the underlying mock observations.

### MCMC

So-called likelihood-free inference (LFI) algorithms, such as rejection ABC, lend themselves to our analysis because the likelihood that describes the system is intractable. By introducing a distance measure based on suitable summary statistics, rejection ABC thus altogether avoids the necessity of consulting the likelihood when inferring the posterior distribution.

Alternatively, one might introduce a surrogate likelihood. This approach is, indeed, commonly used across different fields of research [60, 61].

Following similar arguments to those that lead to the distance measure in Equation 1, we arrive at a surrogate likelihood (*p*(**Θ**|***x***)) in the form

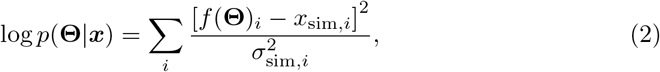

where *x*_sim,*i*_ and *σ*_sim,*i*_ again denote the median and standard deviation of the marginal of the *i*th output variable in the mock observations, respectively.

However, the use of Equation (2) leads to convergence problems, due to the stochastic nature of the ABMs. Alternatively, one might directly compare the predicted median (*M*) with that of the observations. To account for the stochastic behaviour of the ABMs, we include the standard deviation (*σ*_pred_) of the prediction in the denominator:

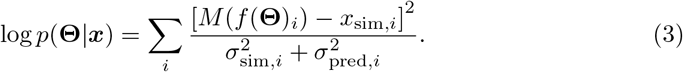

We refer to the surrogate likelihoods in Equations (2) and (3) as case a and b, respectively.

Having defined a surrogate likelihood, we can employ standard sampling techniques, such as Markov chain Monte Carlo (MCMC) [6, 62]. The advantage of this approach over that of rejection ABC is that we can sample more efficiently from the prior. However, as discussed in the section *Results* and mentioned above, the stochasticity of the emulator, partly reflecting the stochasticity of the ABM, might compromise convergence.

In this paper, we use the ensemble sampler published by Foreman-Mackey et al. [63] based on the procedure suggested by Goodman & Weare [64]. We thus evolve an ensemble of *N* walkers in parallel, where *N* is twice the number of dimensions of the mapped input parameter space of the ABM. For each combination of parameters in our mock observations, we exclude a burn-in phase containing 5000 samples per walker. We then collect 20,000 additional samples per walker. To assess the robustness of the MCMC, we estimate the integrated autocorrelation length using an autoregressive-moving-average model. In short, the autocorrelation length denotes the number of samples that is necessary for the walker to become oblivious to its initial conditions. For further details, we refer the interested reader to the paper by Foreman-Mackey et al. [63].

### Direct inference

The procedures presented in the section *Emulators* require two separate steps. First, we train and validate an emulator. Then we impose Bayesian statistics. Alternatively, we can construct an ML algorithm that directly predicts **Θ** given points from the output variable space of the ABMs. In the following, we will refer to such an algorithm as a direct inference machine.

Direct inference machines circumvent subsequent Bayesian sampling schemes, such as ABC or MCMC [12, 13]. Once direct inference machines are trained, they are thus much faster to employ than emulators. We quantify this statement in Fig 5 in the results section. However, this advantage comes at the cost of lower flexibility and reduced transparency. Let’s begin with the latter drawback: It is straightforward to assess the performance of an emulator. We simply need to address its ability to mimic the output of the simulations. Not only can we assess the performance of the emulators through the inferred parameter estimates, but we can directly compare the emulators to the underlying simulations through this intermediate step. Since direct inference machines do not contain such an intermediate step, they do not allow for such comparisons. We must fully rely on performance measures that assess the final predictions for **Θ** (cf. the section *Metrics*). Having established an emulator, it is also straightforward to tweak the priors or alter the likelihood function or distance measure during the inference stage, i.e. using the ABC or MCMC. It is only necessary to re-train the emulator if we extend or change the parameter space that we explore.

For direct inference machines, on the other hand, each such change would require the ML algorithm to be re-trained. It is in this sense that direct inference machines are less flexible than emulators are. Of course, this property only becomes an obstacle for direct inference machines if the training phase is a computational bottleneck, which is not the case for our examples.

Both emulators and direct inference machines thus have their advantages and shortcomings. Rather than resorting to a one-size-fits-all approach, one must hence choose between different algorithms based on the inference problem that one faces. Indeed, both emulators and direct inference machines are very popular in different research fields [13, 21, 46].

In this paper, we compare the results obtained using the emulators with those obtained using two direct inference machines. As our first direct inference technique, we employ a broad, deep neural network. Apart from the fact that the input and output spaces are swapped, the architecture of this neural network matches that described in the section *Neural networks* above. As our second approach, we train a Gaussian process based on the settings summarized in the section *Gaussian Processes*.

Before commencing, we still need to address one question: How do we establish credibility intervals for the direct inference machine? Consider the case, where we repeatedly generate samples using one of our direct inference machines based on the *same* set of singular values for each of the input variables. The chosen direct inference machine would then yield a distribution that reflects the stochastic behaviour of the training data and the limited knowledge of the ML algorithm. However, following this approach, we do not include the uncertainty of the observations. Remember that the input variables of the direct inference machines are the observations, and our mock observations are distributions rather than singular values. We should thus not run the direct inference machine repeatedly using the *same* set of input variables. Rather, to include the observational errors, we would need to generate samples from a distribution of input parameters that corresponds to the distribution of the observations [46]. As discussed in the section *Rejection ABC*, we assume that the marginal distributions of the observed quantities are approximated by independent Gaussian distributions. Following the outline by Bellinger et al. [46], we thus sample 10,000 combinations of input parameters for the direct inference machines using a multivariate Gaussian distribution with a diagonal covariance matrix matching the observational constraints. The means of this multivariate distribution are set to the medians of the observations, while the variance reflects the standard deviations of the marginals.

### Metrics

Due to the stochastic nature of the ABMs, we arrive at distributions for their output variables when repeating runs with the *same* input parameters. To quantify how well our emulators recover these distributions, we call on four measures. For each measure, we compare properties of the marginal distributions predicted by the emulators to those obtained from the simulations, i.e. from the ground truth of the mock observations:

- As our first measure, we compute the error in the predicted means of the marginal distributions, i.e. *µ*_pred_ − *µ*_sim_. Here, the subscripts ‘pred’ and ‘sim’ refer to the predictions by the emulators and the results obtained from simulations, respectively. Any deviations by the mean of this measure from zero would reveal a bias. Since we cover a broad region of the parameter space for each ABM, we cite this measure in units of the standard deviation of the marginal obtained from the simulations (*σ*_sim_).
- As our second measure, we include the relative absolute error in the predicted median, i.e. |*M*_pred_ − *M*_sim_| /*M*_sim_. Like the first measure, this measure quantifies the accuracy of the emulator.
- Thirdly, we compute the ratio between the predicted standard deviation of the marginal distribution and the standard deviation obtained from simulations (*σ*_pred_/*σ*_sim_). This measure should be close to one. Otherwise, the emulator over- or underestimates the uncertainties of the predictions.
- Finally, we compute the Wasserstein distance that quantifies the discrepancy between probability distributions [65]. The lower this measure is, the more similar the predicted distribution by the emulator is to the ground truth.

We note that the metrics chosen for the output of the emulators reflect our assumptions about the nature of the data that we would obtain in a real-world scenario. As elaborated upon in the section *Rejection ABC*, we thus assume the observations to be approximated by a multivariate Gaussian distribution with a diagonal covariance matrix. Although we deem the listed measures to be the most suitable, it is still worth stressing that generalisations of the measures above as well as alternative measures exist [13].

Meanwhile, the assumption that the observations are normally distributed does not affect the metrics, with which we assess the inference results, i.e. the obtained posterior distribution for the model parameters. The marginal posterior distributions might very well be skewed or multi-modal, and the metrics should be able to handle this. We turn to these metrics below.

As regards the ABC runs, the MCMC runs, and the direct inference machines, we can compare the obtained distributions to the original input parameter values (**Θ**_o_) of the mock observations. To assess the performance of the algorithms, we thus compute both the residuals and relative difference between the mean of the predicted marginal distributions and the corresponding elements (Θ_o_) of **Θ**_o_. In addition, we compute

- the negative logarithm of the probability density, *q*(*θ*_*o*_), that is attributed to the true value by the algorithm. The average, −𝔼[log *q*(*θ*_*o*_|***x***_*o*_)], of this measure across input and output variables (Θ_o_, ***x***_*o*_) is widely used in the literature [13, 66]. The lower the value of −log *q*(*θ*_*o*_) is, the better the algorithm is at recovering the ground truth. Note that this metric does not build on any assumptions regarding the shape of the marginal posterior distributions.

These measures quantify the accuracy and precision of the considered inference techniques. By comparing the measures across the different algorithms, we can put the performance of the individual algorithms into perspective.

As an additional comparison, one might propose to infer the model parameters using the actual simulations in tandem with the ABC or MCMC algorithms. We could then use the obtained distributions as benchmarks. However, since we are dealing with complex ABMs, any analysis involving the actual simulations become a herculean task that is beyond the scope of our computational resources. After all, this stumbling block was the reason for turning to emulators and direct inference machines in the first place.

## Acknowledgement

This work was supported by The Oli Hilsdon Foundation through The Brain Tumour Charity, grant number (GN-000595) in connection with the program “Mapping the spatio-temporal heterogeneity of glioblastoma invasion”.

## Brain tumour, CA model

A myriad of complementary methods offer ways to construct spatio-temporal agent-based models for cancer. We kindly refer the interested reader to the review by Liedekerke et al. [5] for a comprehensive overview [67–70].

In agent-based cancer models, each agent represents a single tumour cell or cell cluster. In this paper, we settle for a lattice-based cellular automata (CA) model [25, 71]. This choice implies that each tumour cell is associated with a lattice site. For simplicity, we confine our tumour cells to a two-dimensional plane — as in the case of cell growth *in vitro*, i.e. in a cell-culture dish. To avoid artefacts stemming from the geometry of the lattice, we use an irregular grid (cf. Voronoi tessellation [72]). A Cartesian grid would thus lead to a square tumour. Our lattice contains 40,000 lattice sites and does not allow for cells or nutrients to flow beyond its outer boundary — cells at the boundary simply have no lattice sites to expand into, and we impose zero-flux (Neumann) boundary conditions on the nutrient flow.

To encode the dynamics of the cell population, each cell is governed by a set of deterministic and stochastic rules. Modelling the evolution of the system in discrete time steps, we successively apply these rules to each cell. For each time step, we randomly shuffle the order by which we cycle through the cells to avoid introducing spatial preferences and artefacts in the tumour growth.

At any given time step, each cell might migrate to a neighbouring lattice site, divide, or die. At any given time step, cells might die with probability *P*_cd_. A cell can migrate with a probability *P*_mig_ if there is a free neighbouring lattice site to move to. Moreover, cells are not allowed to migrate if cell division takes place during the same time step (cf. the “Go or Grow” hypothesis for glioma, [73]). Since cell division leads to the creation of new cells, cell division can only happen if there is a lattice site available for the daughter cell [74]. If a cell is placed at the tumour edge, it might divide with probability *P*_div_. Cells that are embedded within the tumour can also divide but will have to push their neighbours aside to do so. Since this effort costs energy, the probability for cell division decreases exponentially with the distance to the tumour edge [75, 76]. Here, we set the probability for cell division to zero if the next available spot is more than 8 lattice sites away. Only cells close to the edge, in the so-called proliferating rim, are hence able to divide. The remaining bulk of the tumour is quiescent. This behaviour is consistent with the well-known linear growth of spherical avascular tumours [77]. We also note that this behaviour makes it superfluous to resolve single cells throughout the tumour. Thus, to lower the computational expense, we only monitor cell clusters when we are not dealing with cells in the proliferating rim, i.e. we employ an adaptive irregular lattice.

Glioblastoma multiforme is hierarchically organised, which implies that the tumour is composed of different cell types [45]. This notion is important since each cell type is governed by different rules involving different probabilities (*P*_mig_, *P*_div_, and *P*_cd_). We must, therefore, distinguish between glioblastoma stem-like cells (GSC), propagating progeny (GPP), and cells in a differentiating subpopulation (GDS). There are two possible outcomes when a GSC undergoes cell division: It either splits into two GSCs with probability 1-*ϵ* or a GSC and a GPP with probability *ϵ*. A GPP, on the other hand, might divide into two GPPs or produce one GDS with equal probability. GDSs do not divide.

The cells inhabit an environment that contains nutrients. Initially, these nutrients are evenly distributed across the lattice. However, we have placed a nutrient source in the lower left-hand corner of the lattice, and the nutrients are locally consumed by the cells, as the tumour grows. Gradients in the nutrient concentration (*n*) thus arise over time. The resulting diffusion of nutrients across the lattice is modelled using a reaction-diffusion equation on the form [78–80]

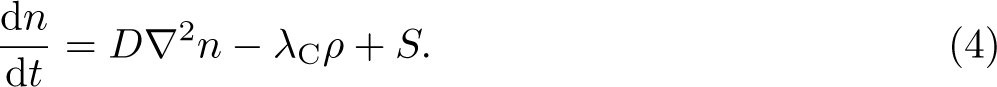

Here, *D* denotes the diffusion coefficient for the nutrient flow, *λ*_C_ scales the nutrient consumption of the cells, and *ρ* denotes the tumour cell concentration, i.e. we assume that all cell types consume the same. Finally, *S* signifies any nutrient sources, i.e the aforementioned inflow of nutrients in the lower left-hand corner. Note that Equation (4) is solved on a Cartesian grid. At each time step, interpolation is hence required between this grid and the irregular lattice on which the cells live. We illustrate the described set-up in Fig 6.

**Fig 6.**
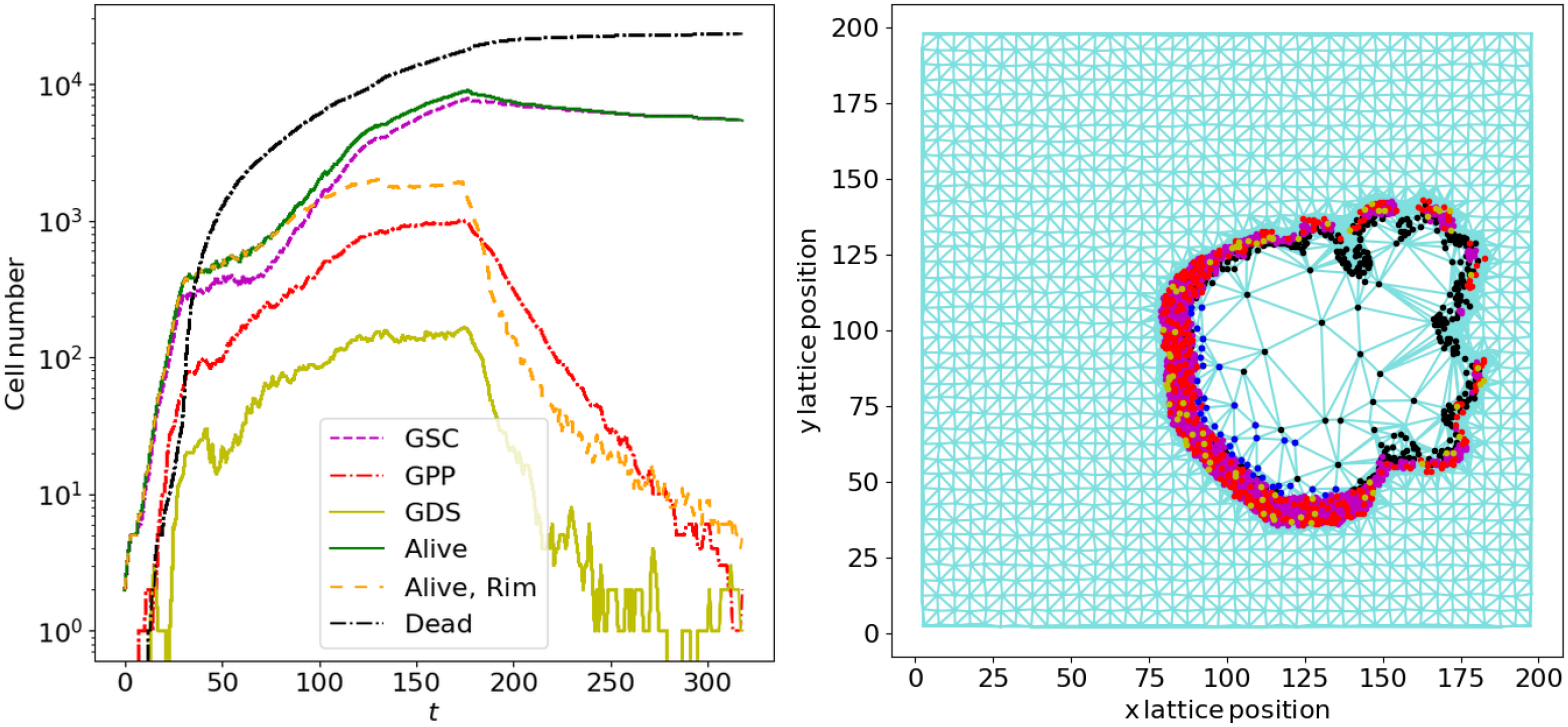
Example summary of the output of the brain cancer CA model for a single model run. The **left panel** summarizes the tumour growths in terms of the size of the different cell subpopulations. For this run, *P*_div_ = 0.25, *P*_cd_ = 0.05, *λ*_C_ = 0.1 units of nutrients consumed per cell per time step, and *ϵ* = 0.2. We include the number of GSC, GPP, GDS, and dead tumour cells as well as the total number of alive tumour cells and the number of alive tumour cells in the proliferating rim. The **right panel** shows a snapshot of the tumour at *t* = 100. The reaction-diffusion equation that describes the nutrient flow is solved on a Cartesian grid. This grid is highlighted in cyan outside the tumour. The cells live on an adaptive irregular grid that can, at most, harbour 40,000 cells. This adaptive grid is highlighted in cyan inside the tumour. In the proliferating rim, the irregular grid has a high resolution. In the bulk tumour, several cells share one lattice site. Single GSC (purple), GPP (red), and GDS (yellow), are shown using markers, whose colours match those in the left panel. Dead tumour cells or clusters, containing only dead cells, are marked with black dots. Clusters with quiescent cells are blue. The presence of a nutrient source in the lower-left corner and the imposed cell death upon starvation introduce an asymmetric growth pattern.

**Fig 7.**
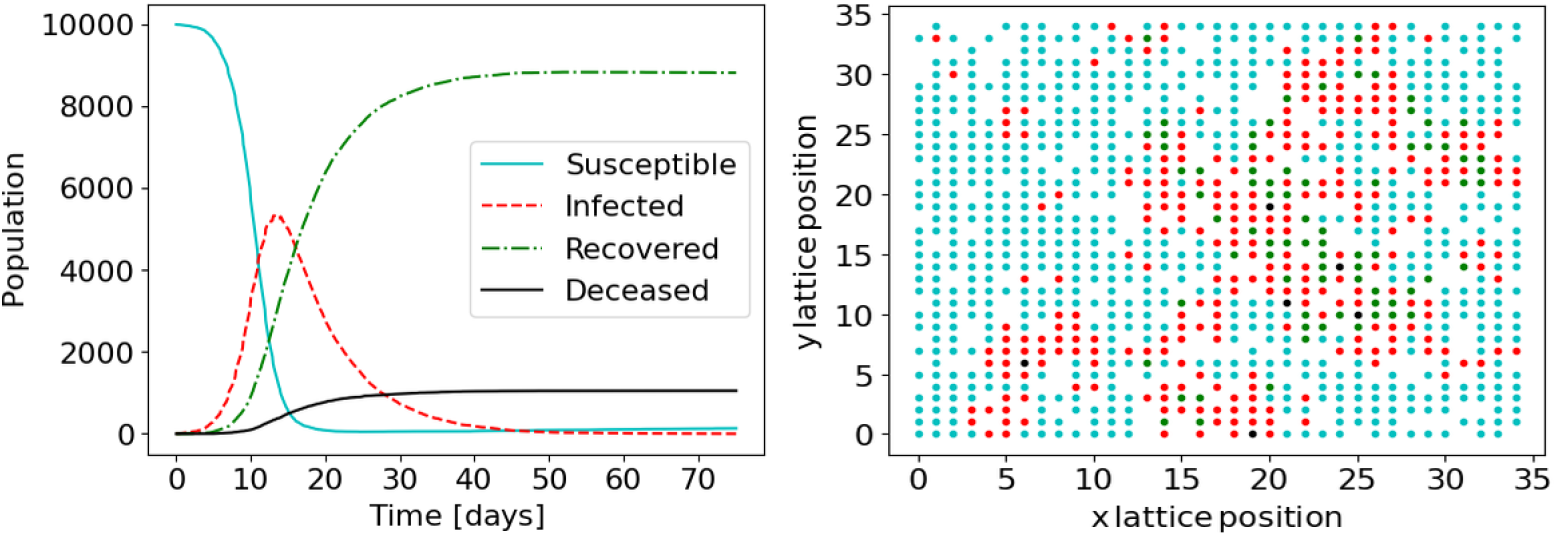
Example summary of the output of the epidemiological SIRDS ABM for a single model run. The **left panel** summarizes the time evolution of the different subpopulations (*S, I, R*, and *D*) with 10,000 individuals randomly distributed over 12,100 lattice sites. In this scenario, *P*_d_ = 0.1, *P*_i_ = 0.2, (*N*_re_) = 5.1 random encounters per infected individual per day, *t*_i_ = 7.1 days, and *t*_p_ = 5000 days. For illustrative purposes, the **right panel** shows a snapshot at *t* = 6.0 days for a simulation with the same parameter values but only 1000 agents randomly distributed over 1225 lattice sites. The colour coding matches that in the left panel: cyan dots mark susceptible individuals, red dots mark infected individuals, recovered individuals are green, and black dots mark dead agents.

We assume that a sufficiently low nutrient concentration will inflict immediate cell death [81]. Such low concentrations might occur at the centre of the tumour leading to a necrotic core. This morphology is observed for real tumours.

For glioblastoma multiforme, the migration of tumour cells is known to primarily occur along white matter tracks and blood vessels [82–84]. One might take this behaviour into account by letting the stochastic rules that govern cell actions vary across the lattice. In addition, one could allow for mutations that alter these rules. However, for our purposes, simple rules for the cell dynamics suffice. Our main focus lies in extracting general guidelines for stochastic models rather than capturing all aspects of this specific brain cancer type.

### Parameters and summary statistics

We initialize each simulation with a single GSC near the centre of the lattice and vary four input parameters (**Θ**):

- Our first parameter is the probability, *P*_div_, for GSCs with a free neighbouring lattice site to divide during each time step. While GPPs are assumed to divide 0.6 times this rate, GDSs do not divide. Note that these rules are chosen for their simplicity rather than given a biological justification.
- The second parameter is the probability, *P*_cd_, for GPPs to undergo programmed cell death during each time step. While GDSs are assumed to be three times more likely to die at every time step, GSCs do not die spontaneously. Again, these rules are chosen for their simplicity. Note that cells die irrespective of their cell type if the nutrient concentration falls below a critical level.
- Our third parameter is *λ*_C_ in Equation (4).
- Finally, we include *ϵ* that denotes the probability for GSCs to differentiate into GPPs during cell division.

Note that *λ*_C_ is the only parameter that is not a probability. However, we choose the values for the remaining parameters (*D* and *S*) in Equation (4) such that interesting scenarios arise when *λ*_C_ is varied between 0.0 and 1.0. While this choice is not biologically motivated, it means that all parameters can be treated in the same way, which provides an interesting contrast to the epidemiological SIRDS model presented in the appendix *Epidemic, SIRDS model*.

We note that our cancer CA model has many more parameters than the ones we vary in our study. Among others, these parameters include *D, S*, and the critical nutrient level below which immediate cell death due to starvation occurs. We set these to 5 units of area per time step, 500 per time step at the lower-left corner, and 0.1 per unit area, respectively. We do also not vary the migration probability of the different cell types. For GSCs and GDSs, we set the probability for cell migration to 0.02, while GPPs migrate with a probability of 0.5. Due to the “Go or Grow” hypothesis, however, the migration probability of any individual cell is temporarily set to zero if the cell has undergone cell division within the same time step.

To decide on the summary statistics, we must consider the type of data that one might encounter in a cancer study. In an animal-based experiment, we might sacrifice the animal to carefully study its brain. Alternatively, we might deal with magnetic resonance imaging (MRI) or in vivo imaging system (IVIS) data. In any case, we would know the time since the injection of the tumour cells into the animal. We hence assume that our data will comprise a detailed snapshot of the tumour rather than a time series for tumour growth. In the best-case scenario, we might have access to, say, spatial transcriptomics (ST) data that enables us to distinguish different cell types and resolve single cells.

The raw cell count already provides extensive information about **Θ**. The reason for this is twofold. First, since we do not vary the initial conditions, variations in **Θ** do not imply significant spatial variation that might affect tumour growth. Secondly, we do not intend to infer spatial properties, such as varying initial conditions or the impact of cell migration. Based on these considerations, we settle for the following five summary statistics:

- The number of alive glioblastoma stem-like cells (GSC) at *t* = 100.
- The number of alive propagating glioblastoma cells (GPP) at *t* = 100.
- The number of alive differentiating glioblastoma cells (GDS) at *t* = 100.
- The number of dead tumour cells at *t* = 100.
- The total number of tumour cells in the proliferating rim at *t* = 100.

As discussed above, we define the proliferating rim as those cells that are placed within 8 lattice sites of a free lattice site. Note that the maximal width of the proliferating rim (*d*_rim_), i.e. here 8 lattice sites, is yet another parameter that we keep fixed in our analysis. Also, note that the probability for cell division is set to decrease for cells that are embedded within the proliferating rim, reflecting the work needed to push other tumour cells aside. For a tumour cell with distance *d* to the nearest free lattice site, the probability for cell division is thus set to be a factor of 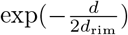 smaller for *d* < *d*_rim_.

## Epidemic, SIRDS model

Agent-based models are in widespread use within the field of mathematical epidemiology. They have thus been applied to a broad variety of data, including studies of the SARS-CoV-2 pandemic [14, 15].

The considered agent-based models for infectious diseases fall into the category of compartmental epidemiological models [85]. Such models, at the very least, distinguish between the infected (*I*) and susceptible (*S*) individuals or subpopulations. In addition, many diseases lead to immunity and some infections might be fatal, making it essential to track recovered (*R*) individuals as well as the death toll (*D*). To indicate that our model takes these four categories into account, we refer to it as a SIRDS model. By repeating the reference to the susceptible individuals (S), we express that recovered individuals might become susceptible to the disease again after a certain period of immunity.

One might model the agents as a set of Brownian particles that move in a two-dimensional plane. Much can already be learned from such an approach and the correct macroscopic behaviour can be reproduced [86]. This being said, human interactions are governed by rules that reflect socio-demographic structures. In addition to random encounters, people thus tend to have predictable encounters within a certain social group. Regular social interactions with the same individuals take place within households, at educational facilities or workplaces. Agent-based models present an opportunity to include such social networks [4, 15, 87–89].

In this paper, we consider agents on a square lattice (see [23, 26, 27, 90, 91] for other examples of lattice-based models). In addition to having *N*_re_ random encounters, each agent interacts with the agents at the eight neighbouring lattice points on a daily basis. Leaving random encounters aside, we note that the agents do not move throughout the simulation, i.e. the social networks are rigid. The distance between agents thus expresses social relationships rather than geographical proximity. In total, there are 12,100 (110 × 110) available lattice sites. To account for the variability of social groups, only 10,000 randomly selected sites are occupied. A susceptible agent that encounters an infected agent will itself be infected with a probability *P*_i_ (for each day of exposure). There are originally three infected individuals randomly scattered across the lattice. An infected individual will recover or die after a number of days drawn from an exponential distribution with mean *t*_i_. The agent dies with the probability *P*_d_. If the agent recovers, it will be protected from being infected during a number of days drawn from an exponential distribution with mean *t*_p_. The agent will then re-enter the population of susceptible individuals.

We stress that our epidemiological SIRDS model is rather simplistic. For instance, factors, such as an individual’s socioeconomic group, sex, or age, might play a role in mortality and infection rates. Many mathematical models thus build on census data to draw a more realistic picture [92, 93]. We do not take such information into account. However, the considered level of complexity is sufficient for our purposes. Rather than modelling a specific disease, our focus lies on the stochasticity that all epidemiological agent-based models share, irrespective of their level of detail.

### Parameters and summary statistics

In the case of our SIRDS model, we vary five input parameters (**Θ**):

- The first parameter is the infection probability, *P*_i_, per day during contact with an infected individual. Since *P*_i_ denotes a probability, this parameter can take values between 0.0 and 1.0.
- Our second parameter is the mortality rate, *P*_d_, of the disease. Again, this parameter can take values between 0.0 and 1.0.
- As a third parameter, we include the time, *t*_i_, during which the infected person can transmit the disease. We evolve the system in time steps, d*t*, of 0.1 days with an agent becoming contagious from the time step that follows the infection. As regards the model grids presented in the section *Model grid*, we vary this parameter on a linear scale from 1.0 to 30.0 days.
- Our fourth parameter is the time, *t*_p_, during which a recovered individual is protected from getting the disease. We vary this parameter on a logarithmic scale between −2.0 and 4.0 when constructing the grids presented in the section *Model grid*. At low values for *t*_p_, the individual does not build up any immunity. Such models are appropriate for some bacterial infections, including meningitis. At high values for *t*_p_, the disease effectively confers lifelong immunity as in the case of some viral diseases, such as measles [94].
- Finally, we vary the number, *N*re, of random encounters with the general population. For the grids in the section *Model grid*, we use a logarithmic scale from −2.0 to 2.0. The model is hereby, for instance, able to account for social distancing and lockdown measures.

In contrast to the brain cancer model, we hence include parameters on different scales.

Rather than being interested in the spatial properties of our specific model, we aim to gain insights into the general properties of agent-based models for infectious diseases. When choosing our summary statistics, we hence turn to the emergent properties of the simulations. These properties comprise the time evolution of the different subpopulations, i.e. *S*(*t*), *I*(*t*), *R*(*t*), and *D*(*t*). To facilitate an easy comparison with our cancer model, we distil these time series into five summary statistics:

- Our first measure is the duration of the outbreak. Note that the computation is interrupted if the duration exceeds 1825 days.
- Secondly, we include the total death toll at the end of the outbreak.
- The third measure is the largest number of infected individuals during the outbreak. Note that we only record the largest peak, although the epidemic might lead to several waves.
- Fourthly, we include the time at which the largest peak in infection numbers occurs. To motivate this choice, let’s assume that the first peak in infection numbers is the largest. The ratio between the time at which it occurs and the total duration of the epidemic would then hint at the existence of multiple waves and help to constrain *t*_p_.
- Finally, we include the largest number of recovered agents during the epidemic. In contrast, the number of recovered people at the end of the epidemic is not a suitable measure because we include scenarios, where the disease does not confer immunity.

